# The dandelion *PARTHENOGENESIS* gene dominantly modifies *Arabidopsis* fertilization and embryogenesis

**DOI:** 10.64898/2026.08.25.747015

**Authors:** Rita B. Lima, Yazhong Wang, Zhihua Cheng, Nicole Jansen, Dilsha Kheani, Violet Sackett, Yannick Jacob, Charles J. Underwood

## Abstract

Parthenogenesis of totipotent egg cells is rare, yet widespread, across the tree of life but mechanistic insights into factors that control parthenogenesis remain sparse. The *Taraxacum officinale PARTHENOGENESIS* (*ToPAR*) gene encodes a C2H2-zinc finger and EAR domain containing protein which is required for parthenogenesis and clonal seed production in apomictic dandelions^1,2^. Ectopic expression of *ToPAR* can trigger egg cell division in lettuce and maternal haploid induction in foxtail millet^1,3^, and *ToPAR* has been employed in a high-penetrance synthetic apomixis system in hybrid rice^4,5^. To date a convenient model system to study *ToPAR* function has yet to be established nor has the capacity for *ToPAR* to trigger cell division in non-gametic cells been tested. Here, we demonstrate that expression of *ToPAR* in egg cells of *Arabidopsis thaliana* using the *EGG-CELL 1.1* promoter (*pAtEC1.1)* causes a reduction in seed set and can trigger egg cell division without fertilization. We found that the *pAtEC1.1:ToPAR* transgene is rarely transmitted through the female lineage where it causes aberrant cell divisions. Expression of *ToPAR* in sexual embryos under the *WUSCHEL RELATED HOMEOBOX 8* (*AtWOX8)* promoter alters cell patterning disrupting morphogenesis. Our results demonstrate that *A. thaliana* can be a powerful system to dissect the mode of action of ToPAR, and that gamete-specific co-factors are not essential for its function.

## Results

To investigate whether *ToPAR* could impact *Arabidopsis thaliana* (Arabidopsis) female reproductive development, we expressed the *Taraxacum officinale PARTHENOGENESIS* (*ToPAR*) gene in Arabidopsis egg cells using the regulatory sequence of the native *EGG-CELL 1.1* (*pAtEC1.1*) promoter^6^. We first screened *pAtEC1.1:ToPAR* T1 plants for fertility defects by quantifying seed phenotypes in near-dry selfed siliques. While wild-type (WT) siliques contained mostly green seeds (green), *pAtEC1.1:ToPAR* siliques additionally carried shriveled-looking seeds (aborted) and occasionally white seeds (white) (**Figure 1A**; **Figure S1**). We also noticed that some non-mature siliques contained aborted ovules. From 42 independent T1 lines, we selected 4 showing different fertility (ranging from 46.9% to 60%) to further investigate these phenotypes in the subsequent T2 generation (**Figure 1A**). To achieve this, we emasculated T2 *pAtEC1.1:ToPAR* buds, pollinated them with WT (to ensure that the observed fertility phenotypes were caused by maternal *ToPAR* expression) and quantified the seed phenotypes 12 days after pollination (DAP) (**Figure 1B-D**). At 12 DAP, all *pAtEC1.1:ToPAR* independent lines had siliques with pale seeds (33.1-42.7% in line 38; 23.1-36.6% in line 39; 35.3-39.9% in line 42; 43-44% in line 43), where they occurred significantly more often compared to WT where they were rarely observed (**Figure 1C-D**). These pale seeds would later collapse and become shriveled (**Figure S2A**). At 12 DAP the frequency of shriveled seeds was significantly higher in individuals from lines 38 and 39, compared to the WT (**Figure 1D**). Curiously, we did not observe any white seeds in T2 lines, except in line 43 when selfed (**Figure S2B)**. We quantified ovule abortion in non-mature siliques by collecting T2 siliques at 6 DAP (**Figure 1B and Figure S2C**). We observed significantly higher ovule abortion in at least one individual per independent line compared to WT, while some T2 siblings did not display this phenotype (**Figure S2C**). For this reason, we showcase the individuals from each independent line separately throughout this manuscript. Furthermore, early aborting seeds were occasionally observed in some lines (**Figure S2C**). Altogether, this data indicates that *ToPAR* expression in egg cells strongly reduces plant fertility.

**Figure 1.**
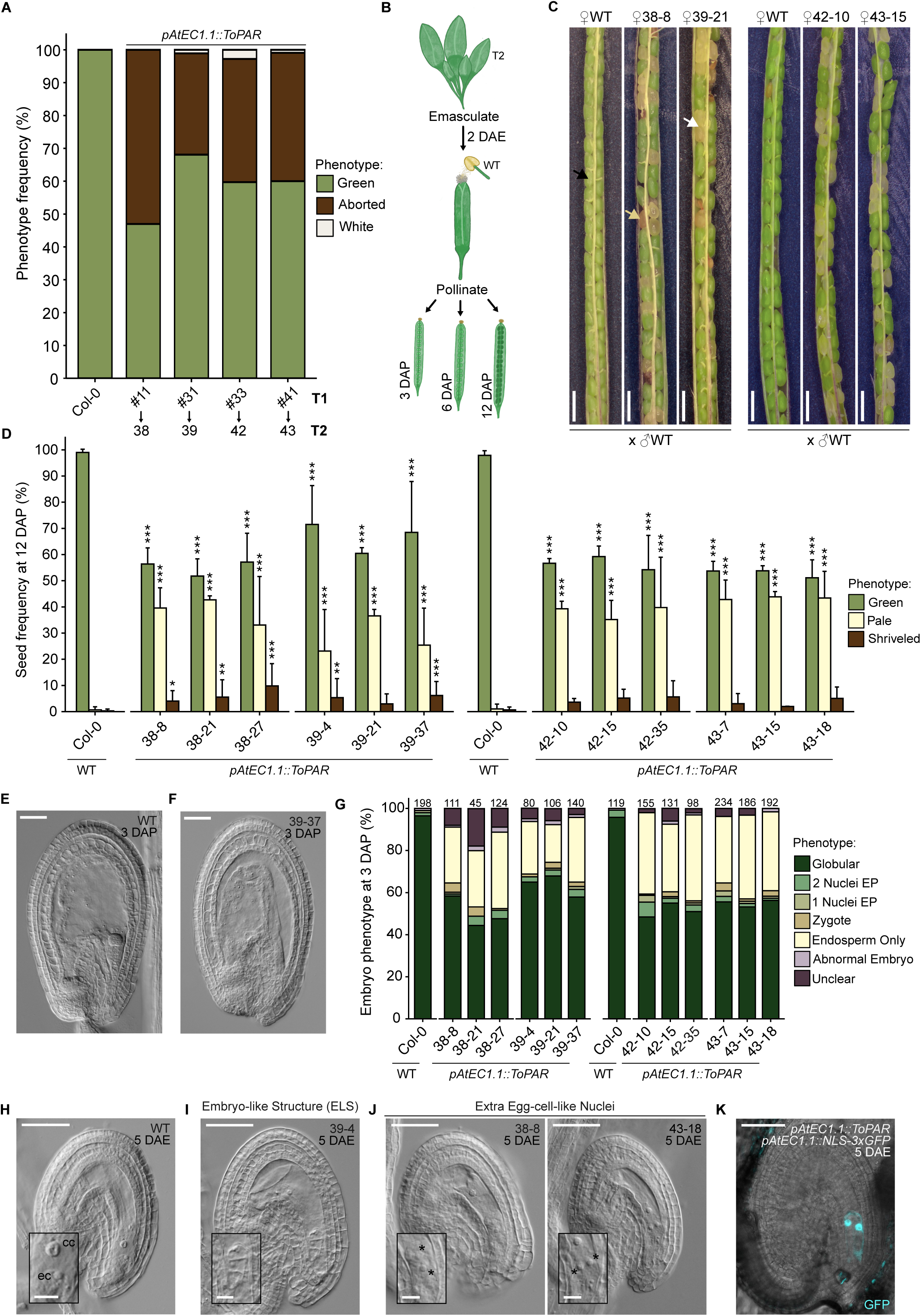
*ToPAR* expression in egg cells reduces seed set and triggers parthenogenesis. (**A**) *pAtEC1.1::ToPAR* T1 seed phenotype screening. Two to three siliques were quantified per independent line. (**B**) Schematic representation of the experimental setup. Buds were emasculated, pollinated 2 days after emasculation (DAE) and siliques collected at 3, 6 and 12 days after pollination (DAP). (**C**) WT and *pAtEC1.1::ToPAR* T2 siliques at 12 DAP. Black, yellow and white arrows denote green, aborted and pale seeds, respectively. Scale bars, 1 mm. (**D**) T2 seed phenotype quantification at 12 days after pollination (DAP). Three siliques were quantified per genotype. Error bars represent standard deviation. Significance of differences was determined by a two-sided Fisher’s exact test. (**E-F**) Microscopic photos of a WT seed with a globular embryo and endosperm (**E**) and of a *pAtEC1.1::ToPAR* seed only containing an endosperm (**F**) at 3 DAP. Scale bars, 50 μm. (**G**) *pAtEC1.1::ToPAR* T2 quantification of embryo phenotype frequency at 3 DAP. Top values indicate the number of seeds analyzed per genotype. (**H-J**) Representative photos of a WT ovule with the visible egg cell (ec) and central cell (cc) (**H**) at 5 DAE and of transgenic ovules showing an embryo-like structure (ELS) (**I**) and extra egg-cell-like nuclei (**J**). Scale bars, 50 μm (12.5 μm for insets). Asterisks denote nuclei in the egg cell apparatus. (**K**) *pAtEC1.1::ToPAR pAtEC1.1::NLS-3xGFP* ovule with two nuclei showing egg cell identity. Scale bar, 50 μm. ****p < 0.0001, ***p < 0.001, **p < 0.01 and *p < 0.05.

We had anticipated that in the T2 generation we would recover 25% homozygotes for the *pAtEC1.1:ToPAR* transgene and that those individuals would have higher rates of unhealthy or aborted seeds, compared to heterozygous siblings. However, from all the T2 individuals analyzed, seed abortion never surpassed 50% (**Figure 1D**). Therefore, we hypothesized that the transgene may not be transmitted via the female lineage. To test this, we pollinated T2 *pAtEC1.1:ToPAR* lines with WT pollen and genotyped the offspring for the presence of the transgene. If female transmission was completely impaired, then one would expect that all *pAtEC1.1:ToPAR* offspring with WT fathers would not carry the transgene. In agreement with rare transmission via the female lineage, the *pAtEC1.1:ToPAR* transgene was only detected in 2 of 419 offspring of transgenic mothers pollinated by WT and average transmission frequencies ranged from 30.4% to 54.3% in selfed offspring, well below the expected 75% (**Table 1**). Curiously, when line 43 was pollinated with WT pollen the white seed phenotype was once again lost, indicating that paternal inheritance of the transgene may be crucial for this phenotype. Consistent with rare transmission of the transgene through the female lineage, ploidy analysis of 210 T3 plants using flow cytometry revealed no haploid offspring (**Figure S2D**).

**Table 1.** *pAtEC1.1::ToPAR* transgene transmission in T2 crossed and selfed offspring.

| Cross | Transgene positive | Transgene negative | Total seedling number | *pAtEC1.1::ToPAR* Transmission (%) |
| --- | --- | --- | --- | --- |
| ♀38 x ♂Col-0 | 0 | 24 | 24 | 0 |
| ♀38 x ♂38 | 40 | 41 | 81 | 49.4 |
| ♀39 x ♂Col-0 | 1 | 120 | 121 | 0.83 |
| ♀39 x ♂39 | 108 | 91 | 199 | 54.3 |
| ♀42 x ♂Col-0 | 0 | 92 | 92 | 0 |
| ♀42 x ♂42 | 35 | 80 | 115 | 30.4 |
| ♀43 x ♂Col-0 | 1 | 181 | 182 | 0.55 |
| ♀43x ♂43 | 86 | 134 | 220 | 39.1 |

To explore the cause of the strong seed abortion, we collected seeds at 3 DAP and analyzed whether embryo development occurred normally (**Figure 1B, E-G**). We hypothesized that egg cell expression of *ToPAR* could impact normal fertilization and/or embryogenesis, thereby causing aberrant seed phenotypes. As expected, WT seeds at 3 DAP showed embryos at the globular developmental stage (95.8-96.5%), which were also seen in the transgenic lines albeit at lower frequency (44.4-58.4% in 38; 57.9-67.9% in 39; 48.4-55% in 42; 53.2-55.6% in 43) (**Figure 1E** and **G**). Strikingly, many *pAtEC1.1:ToPAR* seeds contained what resembled a well-developed endosperm, while lacking an embryo, suggesting that egg cell fertilization is impaired in the transgenic lines (0% in WT; 26.7-36.3% in 38; 17.9-30.7% in 39; 32.1-40.8% in 42; 31.6-39.8% in 43) (**Figure 1F** and **1G**). Additional phenotypes were observed, including embryos at the zygotic stage, embryos with 1 or 2 nuclei at the embryo proper and embryos with abnormal development (**Figure 1G**, **Figure S2E-H**). Moreover, in some seeds it was not possible to distinguish the presence or absence of an embryo (classified as ‘unclear’) (**Figure 1G**). These seeds contained a mass of nuclei in the micropylar region, which could be very dense endosperm or possibly an embryo lacking structure (**Figure S2I**).

The absence of egg cell fertilization could be potentially explained by two mechanisms: (1) the egg cell has lost its gametic potential and/or (2) due to *ToPAR* expression, the egg cell emits a signal that it has been fertilized, thus blocking sperm nuclei fusion. We hypothesized that if the egg cell had lost its gametic potential, then expression of known egg cell identity markers, such as *EC1.1*, would be abolished in lines harboring the *pAtEC1.1:ToPAR* transgene. To test this, we crossed the reporter line *pAtEC1.1::NLS-3xGFP*^6^ *with pAtEC1.1:ToPAR* to generate *pAtEC1.1::NLS-3xGFP pAtEC1.1:ToPAR* and checked for GFP expression in egg cells. Similarly to the *pAtEC1.1::NLS-3xGFP* reporter, fluorescent signals were detected in *pAtEC1.1::NLS-3xGFP pAtEC1.1:ToPAR* egg cells, revealing that despite lack of fertilization, *pAtEC1.1:ToPAR* egg cells have gametic potential (**Figure S2J-L**).

We speculated that if *ToPAR*-expressing egg cells emitted a signal that they were fertilized, then most ovules would attract a single pollen tube for central cell fertilization which appeared normal. On the other hand, if transgenic egg cells were still available for fertilization, then multiple pollen tubes would be attracted to the same ovule to attempt egg cell fertilization. To investigate this, we scored pollen tube number at 12 hours after pollination (12 HAP) with aniline blue staining where WT and *pAtEC1.1:ToPAR* plants were crossed with WT pollen. Multiple pollen tubes (polytubey) entering the ovules were observed in control and *pAtEC1.1:ToPAR* crosses with WT at a low rate, however only one out of ten lines was significantly different compared to the WT (p=0.001) (**Figure S2M** and **N**). These observations suggest that unfertilized *pAtEC1.1:ToPAR* egg cells do not cause additional pollen tube attraction, and that the transgenic gametes present themselves as ‘post-fertilization’ to pollen tubes.

It was previously demonstrated that *ToPAR* expression in egg cells can trigger embryogenesis without fertilization^1,5,7^. Therefore, we questioned if *ToPAR* parthenogenetic potential could also extend to Arabidopsis. To assess this, we collected *pAtEC1.1:ToPAR* T2 pistils 5 days after emasculation (DAE) and quantified parthenogenesis events. An event was only considered parthenogenetic when an embryo-like structure (ELS) with two or more nuclei separated by cell walls could be distinguished within the egg apparatus. We observed the presence of ELS in 3 out of 4 independent lines with frequencies ranging from 1 to 2.4%, which were not detected in any WT ovules analyzed (**Figure 1H-I**, **Table 2**). In addition to ELS, other phenotypes were observed in the transgenic lines and quantified. While in WT embryo sacs a single egg cell and central cell were observed, in *pAtEC1.1:ToPAR* gametophytes multinucleate phenotypes were detected. These included nuclei that resembled extra egg-cells and were often positioned under or next to the egg cell (1-6.5%), additional nuclei that were smaller than egg cells (1-10%) and endosperm-like nuclei (2.2-30.9%) (**Figure 1I, J** and **K**; **Figure S2O, S2P** and **S2Q**; **Table 2**). Strikingly, analysis of *pAtEC1.1::NLS-3xGFP pAtEC1.1:ToPAR* ovules revealed that additional nuclei, besides the egg cell, had acquired egg cell identity as defined by expression of the *pAtEC1.1::NLS-3xGFP* transgene that is usually restricted to egg cells (**Figure 1K** and **Figure S2Q**). These nuclei were either next to the egg cell or where the synergids are located, indicating that they likely arise from egg cell division and/or other embryo sac cells acquiring a new identity. Further phenotypes that did not fit any of the previously mentioned categories were classified as ‘others’ (**Figure S2R**). In conclusion, this data demonstrates that *ToPAR* expression in Arabidopsis egg cells can trigger parthenogenesis, erratic cell divisions and potentially changes in cell identity in female gametophytes.

**Table 2.**
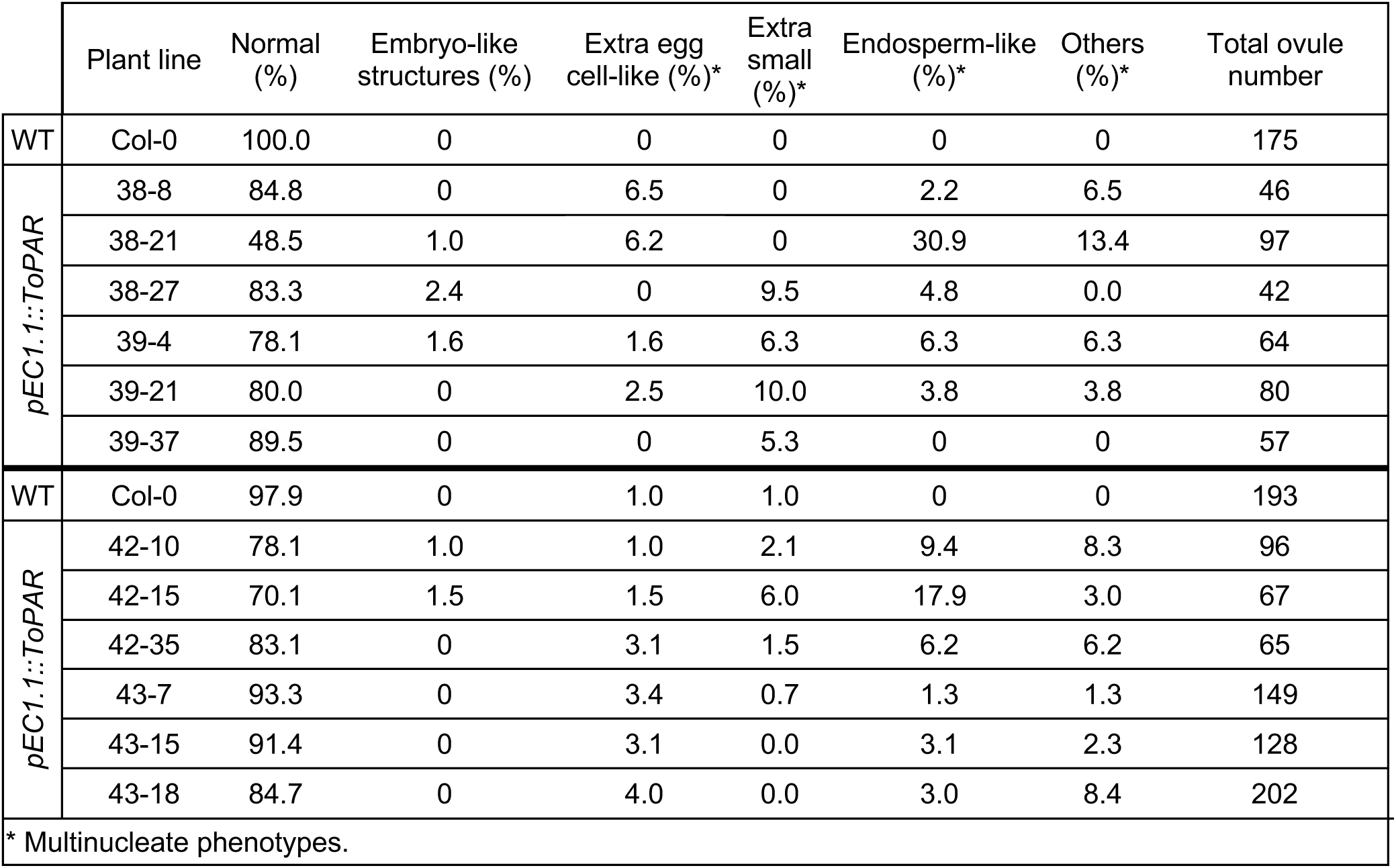
Ovule phenotype quantifications at 5 days after emasculation (DAE).

|  | Plant line | Normal (%) | Embryo-like structures (%) | Extra egg cell-like (%)* | Extra small (%)* | Endosperm-like (%)* | Others (%)* | Total ovule number |
| --- | --- | --- | --- | --- | --- | --- | --- | --- |
| WT | Col-0 | 100.0 | 0 | 0 | 0 | 0 | 0 | 175 |
| *pEC1.1::ToPAR* | 38-8 | 84.8 | 0 | 6.5 | 0 | 2.2 | 6.5 | 46 |
|  | 38-21 | 48.5 | 1.0 | 6.2 | 0 | 30.9 | 13.4 | 97 |
|  | 38-27 | 83.3 | 2.4 | 0 | 9.5 | 4.8 | 0.0 | 42 |
|  | 39-4 | 78.1 | 1.6 | 1.6 | 6.3 | 6.3 | 6.3 | 64 |
|  | 39-21 | 80.0 | 0 | 2.5 | 10.0 | 3.8 | 3.8 | 80 |
|  | 39-37 | 89.5 | 0 | 0 | 5.3 | 0 | 0 | 57 |
| WT | Col-0 | 97.9 | 0 | 1.0 | 1.0 | 0 | 0 | 193 |
| *pEC1.1::ToPAR* | 42-10 | 78.1 | 1.0 | 1.0 | 2.1 | 9.4 | 8.3 | 96 |
|  | 42-15 | 70.1 | 1.5 | 1.5 | 6.0 | 17.9 | 3.0 | 67 |
|  | 42-35 | 83.1 | 0 | 3.1 | 1.5 | 6.2 | 6.2 | 65 |
|  | 43-7 | 93.3 | 0 | 3.4 | 0.7 | 1.3 | 1.3 | 149 |
|  | 43-15 | 91.4 | 0 | 3.1 | 0.0 | 3.1 | 2.3 | 128 |
|  | 43-18 | 84.7 | 0 | 4.0 | 0.0 | 3.0 | 8.4 | 202 |

\* Multinucleate phenotypes.

Ectopic expression of parthenogenetic factors for synthetic apomixis strongly relies on the use of specific promoters that show activity exclusively in egg cells prior to fertilization and early embryogenesis after. Given the low availability of such regulatory sequences, the *pAtEC1.1* and other related variants (e.g. *pAtEC1.2* and *pAtEC1.2en-pAtEC1.1*), are still the most widely used promoters to induce embryogenesis without fertilization. We sought to test the promoter sequence of the Arabidopsis *WOX8* which is specifically active in egg cells and during early embryo development to drive *ToPAR* expression^8^. We generated *pAtWOX8:ToPAR* transgenic lines and followed the same experimental strategy as described above. Out of 15 T1 independent lines screened for reduced fertility, three were selected for more detailed experiments (**Figure S3**, **Figure S4A**). From T2 plants pollinated with WT, siliques were collected 12 DAP and the phenotypes of the seeds were quantified. Similarly to *pAtEC1:ToPAR* lines, we observed the presence of pale seeds that would later collapse in all three independent lines (**Figure 2A**, **Figure S4B**). The frequency of already shriveled seeds was also significantly increased in most individuals compared to the WT (**Figure 2A**). Interestingly, at 6 DAP, we did not identify differences in ovule abortion in comparison to the WT, revealing that *pAtWOX8* driven expression of *ToPAR* is not detrimental for ovule development (**Figure S4C**). In summary, this data reveals that *ToPAR* expression driven by *pAtWOX8* negatively impacts plant fertility.

**Figure 2.**
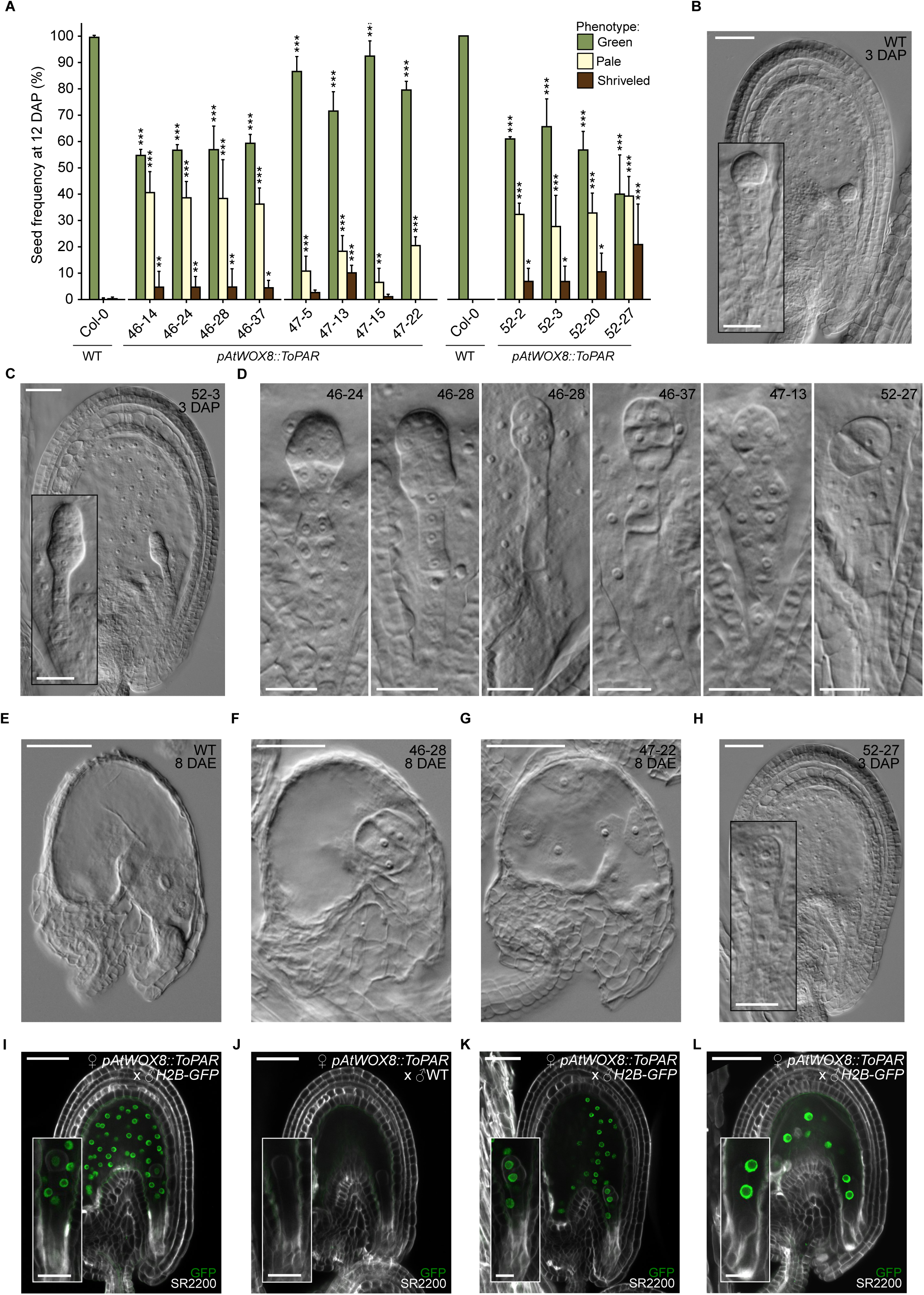
*ToPAR* expression during embryogenesis alters cell patterning and embryo morphogenesis. (**A**) T2 seed phenotype quantification at 12 days after pollination (DAP). Three siliques were quantified per genotype. Error bars represent standard deviation. Significance of differences was determined by a two-sided Fisher’s exact test. (**B-C**) Microscopic photos of a WT seed with a globular embryo (**B**) and a *pAtWOX8::ToPAR* seed with an abnormal embryo (**C**) at 3 DAP. Scale bars, 50 μm. (**D**) Compilation of embryos with abnormal patterning observed in *pAtWOX8::ToPAR* seeds at 3 DAP. Scale bars, 25 μm. (**E-G**) Microscopic photos of a WT ovule (**E**) and *pAtWOX8::ToPAR* ovules with an embryo-like structure (ELS) (**F**) and an autonomous endosperm at 8 days after emasculation (DAE) (**G**). Scale bars, 50 μm. (**H**) *pAtWOX8::ToPAR* seed with an embryo containing one nucleus at the embryo proper surrounded by a more developed endosperm at 3 DAP. Scale bars, 50 μm (25 μm for inset). (**I-J**) Confocal images of seeds resultant from *pAtWOX8::ToPAR* ovules pollinated with *pRPS5A::H2B-GFP* (*H2B-GFP*) (**I**) or WT (**J**) at 2 DAP. Seeds were stained with SR2200 (in white). Scale bars, 50 μm (25 μm for insets). (**K-L**) Confocal images of *pAtWOX8::ToPAR* ovules pollinated with *H2B-GFP* showing a GFP positive abnormal embryo (**K**) and a non-fluorescent embryo surrounded by an endosperm with GFP signal (**L**) at 2 DAP. Scale bars, 50 μm (25 μm for insets). ****p < 0.0001, ***p < 0.001, **p < 0.01 and *p < 0.05.

To determine the cause of seed abortion, we imaged 3 DAP seeds and quantified their phenotypes. We found that the most striking difference between *pAtWOX8:ToPAR* and WT seeds was an increase in embryos with aberrant embryo patterning (1-1.3% in WT; 17.1-22.9% in 46; 5.4-12.3% in 47; 16.3-38.2% in 52) (**Figure 2B-D**, **Figure S4D**). We additionally observed an increase in the frequency of embryos with one nucleus at the embryo proper compared to the WT (0.5-1.9% in WT; 11-14.1% in line 46; 3.9-12.3% in line 47; 5.1-10.9% in line 52) (**Figure S4D**). The abnormalities in embryo patterning were not uniform among seeds and included extra divisions in the suspensor and/or apical cell, extra-long suspensor cells and wrong cell wall orientation (**Figure 2D**). In some instances, it looked as if a second embryo proper was originating in suspensor cells (**Figure 2C and D**). The frequency of the diverse phenotypes also varied among siblings. This data shows that *ToPAR* expression strongly modifies embryogenesis in Arabidopsis.

Next, we tested whether *pAtWOX8:ToPAR* egg cells could initiate parthenogenesis. We did not found evidence of parthenogenesis nor of abnormal embryo sac formation at 5 DAE (data not shown). Since autonomous embryogenesis could initiate later in these lines, we analyzed ovules at 8 DAE. Despite most ovules being aborted at this time point, those that kept their shape could contain normal egg and central cells (classified as ‘normal’) (**Figure 2E**, **Figure S4E**, **Table S1**). Remarkably, ELS structures were found in all independent lines with frequencies between 0.3 and 1.7% (**Figure 2F**, **Table S1**). Curiously, autonomous endosperm was observed both in transgenic and WT embryo sacs (**Figure 2G**, **Table S1**), which is likely related to the age of the non-fertilized ovule. This data supports that *ToPAR* expression in egg cells can trigger parthenogenesis in Arabidopsis.

When quantifying *pAtWOX8:ToPAR* 3 DAP seeds (**Figure S4D**), we noticed that the endosperm of seeds with one nucleus at the embryo proper were often more developed than expected for that embryonic developmental stage (**Figure 2H**). Since these lines were capable of parthenogenesis, we questioned if embryogenesis could be delayed in those seeds, due to a potential parthenogenetic origin of those embryos. We reasoned that the presence of an endosperm could provide extra support to promote parthenogenetic embryogenesis compared to when central cell fertilization does not take place. To test this, we pollinated *pAtWOX8:ToPAR* pistils with *pRPS5ɑ::H2B-GFP*^9^ (*H2B-GFP*) pollen and screened for GFP expression in the embryos and endosperms of 2 DAP cleared seeds. If the embryos developed through parthenogenesis, one would expect to only detect fluorescence in the endosperm. We observed fluorescent signals in the endosperms and young embryos of both WT and *pAtWOX8:ToPAR* seeds fertilized by *H2B-GFP* pollen, which were not detected when WT was used as pollen donor (**Figure 2I** and **J**, **Figure S4F** and **G**). These observations indicate that the embryos with one nucleus at the embryo proper are a product of fertilization and that they likely develop slower and/or arrest in development. We also observed GFP signals in abnormally developing embryos, confirming that they also result from fertilization (**Figure 2K**). Moreover, we observed two-celled embryos showing what looked like a symmetric first division rather than an asymmetric one (**Figure S4H**). Given that the first asymmetric division establishes the apical and basal cell fates, it is possible that in some *pAtWOX8:ToPAR* embryos apical-basal fate is not properly determined, thereby leading to abnormal embryo development (**Figure 2D**). Despite fluorescent signals being detected in almost all tested embryos, we found a single seed in which GFP was visible in the endosperm, but absent from the embryo (**Figure 2L**, **Figure S4I**). This would be consistent with an embryo of parthenogenetic origin, as was observed in emasculated *pAtWOX8:ToPAR* plants (**Figure 2F**).

Overall, these results show that *ToPAR* expression driven by *AtWOX8* regulatory sequences reduces plant fertility mostly by causing abnormal embryo development and can occasionally trigger egg cell division without fertilization.

## Discussion

Here, we found that *ToPAR* expression under the regulatory sequences of *AtEC1.1* and *AtWOX8* highly impacts Arabidopsis seed production (**Figure 1A**; **Figure S1; Figure S3; Figure S4A**). In heterozygous *pAtEC1.1:ToPAR* plants, the transgene potently modifies female reproduction to the extent that most ovules that inherit the transgene undergo aberrant reproduction (**Figure 1G**), and the transgene is not transmitted by the female lineage (**Table 1**). Despite this we found that *ToPAR* expression in both transgenic lines can induce egg cell division, albeit at low frequencies and without viable haploid offspring formation. Egg cell division was previously reported at low frequency in Arabidopsis, when expression of *Brassica napus BABY BOOM* (*BnBBM*) was induced under the *EC1.2en-EC1.1* promoter^10^. The low frequency of parthenogenesis was attributed to high *BnBBM* expression, and it could be increased by controlled *BnBBM* induction^10^. Likewise, the low frequency of parthenogenesis by *ToPAR* we reported here may be due to incorrect expression level, timing or location during Arabidopsis female reproductive development, or alternatively that additional factors are required. Remarkably, *ToPAR* expression led to embryo sacs with more than one nucleus with egg cell identity (**Figure 1K** and **Figure S2Q**). Based on their embryo sac placement, these egg(-like) cell nuclei might derive not only from egg cell divisions, but also from leaky *AtEC1.1* expression in the synergids, as has been previously reported^11^, suggesting that the timing and domain of *ToPAR* expression requires fine-tuning. It has recently been demonstrated that the combined expression of the *Oryza sativa BBM1* (*OsBBM1*) and *WUSCHEL-LIKE HOMEODOMAIN 9* (*OsWOX9*) in egg cells strongly enhanced parthenogenesis in rice^12^. However, possible synergistic effects of parthenogenetic factors are yet to be tested in Arabidopsis. Our work reveals that additional factors to *BnBBM* can promote egg cell division in Arabidopsis. Strikingly, using the regulatory sequences of *AtWOX8* to drive *ToPAR* expression we induced changes in embryo patterning and shape, showing that ToPAR can alter cell division patterns during early plant development. This revealed, for the first time in any plant species, that gametic cell identity is not required for ToPAR to trigger cell division. We believe our results are a foundation for using Arabidopsis as a model system to understand the molecular pathway of ToPAR. Taking advantage of cotyledon, root and/or leaf primordia promoters to stimulate ToPAR production could be a powerful tool to unveil the molecular mechanisms of ToPAR. Overall, our results demonstrate that *ToPAR* expression in Arabidopsis gametic and sporophytic cells can modulate cell division, development and morphology.

## Resource Availability

### Lead Contact

Further information and material requests should be addressed to the corresponding author, Charles J. Underwood.

### Materials Availability

Generated plant materials can be requested by contacting the corresponding author, Charles J. Underwood.

### Data Availability

All data is available in the supplementary information.

## Acknowledgments

We thank Prof. Stefanie Sprunck for providing the *pEC1.1::NLS-3xGFP* line and Prof. Tetsuya Higashiyama and Prof. Daisuke Kurihara for the *pRPS5a::H2B-GFP* reporter. The recombinant plasmid *pCAMBIA1305.4* was kindly provided by Prof. Xiaolan Zhang (China Agricultural University). We also thank Sterre van Tol for the technical assistance, Dr. Tobias de Werk for statistical analysis support and Koos Janssen, Walter Hendrickx, Peter Arns, Dorine Schonhoven-van Eeuwijk and Noa Spoor for the plant care. This work was supported by the Radboud University (Crop Biotechnology and Engineering funding from the Executive Board of Radboud University to C.J.U.) and the European Research Council (ERC starting grant 101076355, ‘AsexualEmbryo’ to C.J.U.). This work was also made possible by a grant (R35GM128661) from the National Institutes of Health to Y.J.

## Author Contributions

R.B.L. and C.J.U. designed the experiments. Y.W. and Z.C. developed the *pAtEC1.1::ToPAR* construct. V.S. and Y.J. performed the initial fertility screening. R.B.L., N.J. and D.K. performed the experiments. R.B.L. analyzed the data. R.B.L. and C.J.U. wrote the manuscript with input from all co-authors.

## Star Methods

### Plant Materials

The *Arabidopsis thaliana* wild-type ecotype Columbia-0 (Col-0) was used in this study, as well as the previously published reporter lines *pEC1.1::NLS-3xGFP*^6^ and *pRPS5a::H2B-GFP*^9^. Seeds were sterilized by a 12-minute wash with 70% ethanol supplement with 0.05% Tween-20, followed by two washes with 100% ethanol. The seeds were air dried and plated on ½ MS medium (0.43% MS Basal Salt Mixture, 0.05% MES, 1% Sucrose and 0.8% Plant Agar). The seeds were stratified for 48h at 4°C and transferred to a growth chamber (22 °C; 16 h light/8h dark; 100 µmol s−1 m−2; 70% humidity). If required, the MS medium was supplemented with antibiotics (50 μg/uL kanamycin; 20 mg/μL hygromycin). Plates containing transgenic seeds for hygromycin selection were placed for 6h under light upon stratification and then covered with aluminum foil during 48h. After this period, the plates were moved to continuous light for 48h and lastly placed back into normal light conditions (16h light/8h dark) until ready for soil transfer. The seedlings were transferred to soil after 7-10 days and grown in a climate room (21 °C; 16 h light/8h dark; 150 µmol s−1 m−2; 50% humidity).

### Molecular Cloning and Transgenic Generation

The *pAtEC1.1::ToPAR* construct was generated using the ClonExpress MultiS One Step Cloning Kit (Vazyme, Nanjing, China). To achieve this, the promoter regions of the Arabidopsis *EC1.1* (AT1G76750), the coding DNA sequence (CDS) and 3’ untranslated region (3’ UTR) of the dandelion *ToPAR* gene were amplified from Col-0 and dandelion Tara3 genomic DNA (gDNA) and assembled into the destination vector pCAMBIA1305.4^14,15^ (pre-digested with SalI and BstEII). The construct *pAtWOX8::ToPAR* was generated by Golden Gate technology in two steps^16^. First level-0 (L0) vectors were generated by cut and ligate reactions using BpiI restriction enzyme and T4 DNA ligase (New England Biosciences). The *AtWOX8* (AT5G45980) promoter regions were amplified using Col-0 gDNA as template and cloned into vector pICH41295. Both ToPAR CDS and 3’UTR together with the NOS terminator were amplified from the *pAtEC1.1::ToPAR* construct and assembled into the plasmids pICH41308 and pICH41276, respectively. Before L0 assembly, the ToPAR CDS was domesticated. The final product was then assembled by mixing all L0 plasmids with the level-2 (L2) pICH86966 to cut and ligate via BsaI restriction enzyme and T4 DNA ligase. L0 and final constructs were validated by restriction digestion and whole-plasmid sequencing. All primer sequences and assembly parts are listed in Table S2.

The constructs were transformed into *Agrobacterium tumefaciens* strain GV3101 and then into Arabidopsis thaliana by floral dip^17^. Positive transformants were selected using appropriate antibiotics and validated by genotyping. Genotyping details can be found in Table S2.

### Developmental Assays

Flowers were emasculated at stages 12-13 for all developmental assays^18^. For pollination assays, pistils were hand-pollinated 2 days after emasculation (DAE) and siliques opened at 6 and 12 days after pollination (DAP) to quantify ovule abortion and seed abortion, respectively. For seed developmental analysis, siliques were collected at 3 DAP. *In vitro* pollen tube growth assays were achieved by hand-pollinating pistils 2 DAE and collecting them 12 hours after pollination (HAP). Parthenogenesis was tested by collecting pistils 5 and/or 8 DAE. Reporter imaging was performed at 2 DAP for endosperms and embryos and 5 DAE for egg cells.

### Histological and Fluorescence Analysis

Pistil and silique clearings were performed by fixing the samples in ethanol:acetic acid (9:1) during 24-48h, followed by 3 washes of 10 minutes each with 90% ethanol and an additional 10-minute wash with 70% ethanol. The samples were then incubated in a chloralhydrate solution (66.7% chloralhydrate (w/w), 8.3% glycerol (w/w), pH 5.2) for 48h. The ovules/seeds were separated from the carpels and mounted in a glass slide containing a generous drop of chloralhydrate solution, under the stereomicroscope. The samples were observed using differential interference contrast (DIC) optics in the Leica DM6B microscope complemented with a Leica DMC4500 camera.

Pollen tube staining was accomplished by fixing siliques in ethanol:acetic acid (9:1) for 24-48h, followed by 3x five-minute washes with water and overnight incubation in 8M NaOH. The samples were then washed with water for one hour three times and incubated overnight in 0.1% (w/) decolorized aniline blue at 4°C. The samples were mounted by removing the carpels under the stereomicroscope in a glass slide with a drop of aniline blue solution. The pollen tubes were visualized with the Leica DM2500 microscope and images were captured on a DCF7000T Leica camera. Phenotype frequencies were calculated as the proportion of single tube or polytubey ovules among all scored ovules for each line, with 95% Wilson confidence intervals.

ClearSee alpha seed clearings were performed as previously described by Attuluri and colleagues, but with minor modifications^19^. Opened siliques were initially collected in PBS-T (1x PBS supplemented with 0.1% Triton-X100) and then incubated in fixative (4% Paraformaldehyde in PBS supplemented with 0.1% Triton-X100) for 30 minutes under vacuum infiltration. The fixation process was performed on ice and in tubes covered with aluminum foil for light protection from the fixation step to the staining. The samples were then incubated at 4 °C overnight and, in the next day, the samples were washed three times, 1h each wash, in 1x PBS-T at room temperature. The fixed material was incubated in ClearSee solution (10% xylitol (w/v), 15% sodium deoxycholate (w/v), 25% urea (w/v) in water) supplemented with 100 mM sodium sulfite for 20-30 days under agitation. The clearing solution was changed every day. For this, a stock solution of Clearsee was prepared and the sodium sulfite added freshly to an aliquot. The samples were stained overnight with 0.2% (v/v) Renaissance SR2200 in Clearsee alpha solution. The following day, the staining solution was replaced by Clearsee alpha and incubated for 1h. The samples were dissected in a glass slide with Clearsee alpha to expose the ovules/seeds. This was performed under the stereomicroscope. Reporter live-cell imaging was achieved by mounting ovules in a glass slide with 0.1 mg/mL of propidium iodide (PI). The samples were imaged with the Leica SP8x confocal laser scanning microscope equipped with a 405 nm diode laser, a pulsed white light laser and HyDs detectors. Fluorescence detection was done as follows (in nm; excitation - ex and emission - em): SR2200 - ex 405, em 420-500, GFP - ex 488, em 499-525, and PI - ex 488/514, em 635-719.

### Female Transmission Assays

A grid with 42 squares was hand-drawn on MS media plates and individual seeds were sown per square. For crosses with WT, seeds from two to three siliques were sown, while for selfed seeds two plates were sown per line. gDNA was extracted from leaves of seven to 13- day-old-seedlings and used as a template for transgene detection. Genotyping details are listed in Table S2. Offspring of two to three siblings were genotyped per line and pooled together to calculate transgene transmission per independent line.

### Flow Cytometry

Seeds were individually sown on gridded plates as described above. The ploidy was measured by harvesting and chopping leaves from two-week-old seedlings with razor blades in 1 mL of cold Galbraith’s buffer (45 mM MgCl_2_, 30 mM sodium citrate, 20 mM MOPS, 0.1% (v/v) Triton X-100, pH 7.0) supplemented with RNAse (50 µg/mL). The suspension was filtered through a 0.5 µm filter and dyed with PI (1 mg/mL). The ploidy was determined using the BD FACSMelody (BD FACSMelody™). The flow cytometry results were analyzed using the web tool floreada.io and R Studio (https://www.r-studio.com).

### Statistical Analysis

R Studio was used for data analysis. For 6 and 12 DAP experiments, significant differences were determined by comparing each seed phenotype to Col-0 using a two-sided Fisher’s exact test. P values were adjusted across all comparisons using the Bonferroni method.

For pollen tube staining with aniline blue differences among lines were assessed using a Pearson chi-square test, followed by pairwise comparisons of each line with Col-0 using Fisher’s exact tests. P values were adjusted for multiple comparisons using the Holm method. Samples were considered statistically different when p < 0.05. ****p < 0.0001, ***p < 0.001, **p < 0.01 and *p < 0.05.

---

**Figure S1.**
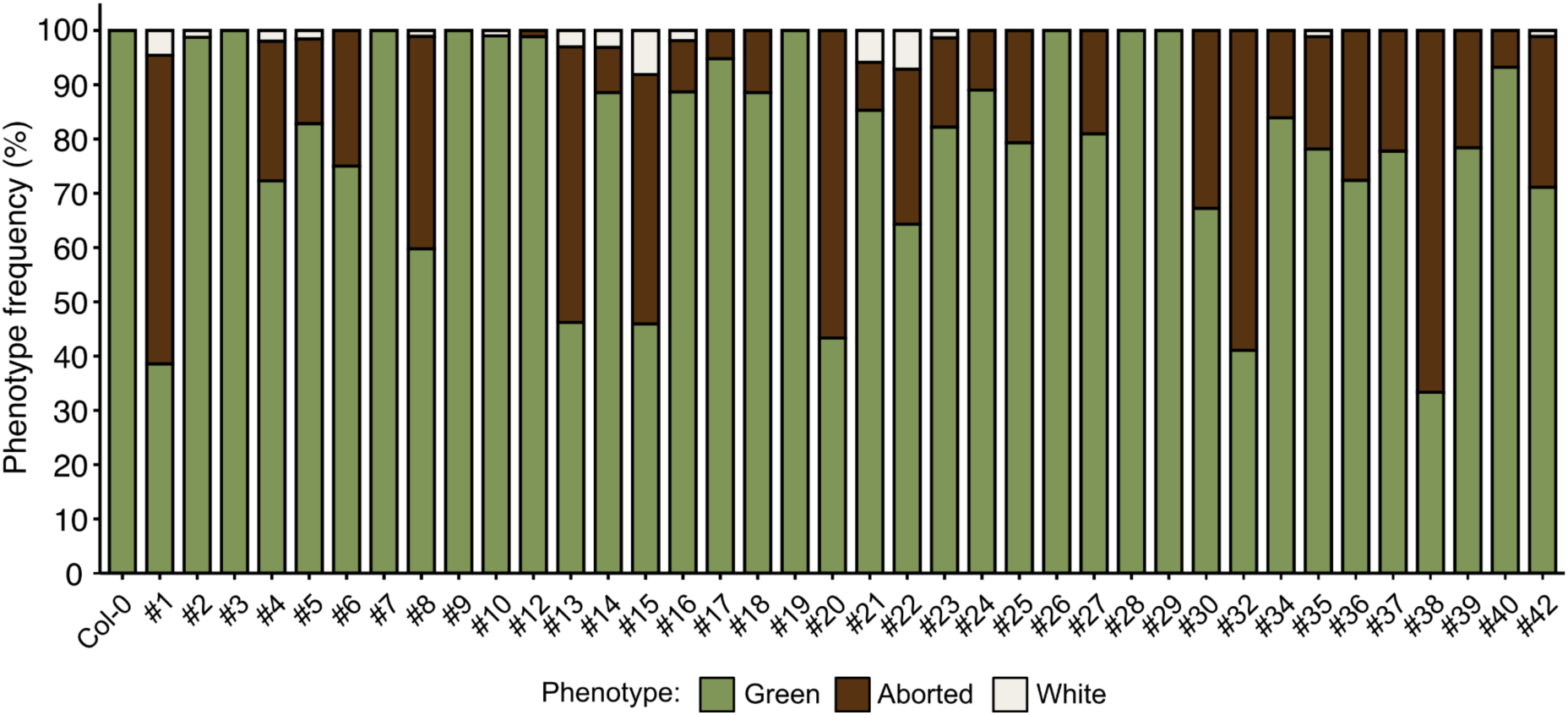
*pAtEC1.1::ToPAR* T1 seed phenotype screening. Phenotype frequencies of additional T1 independent lines. Two-three siliques were quantified per independent line.

**Figure S2.**
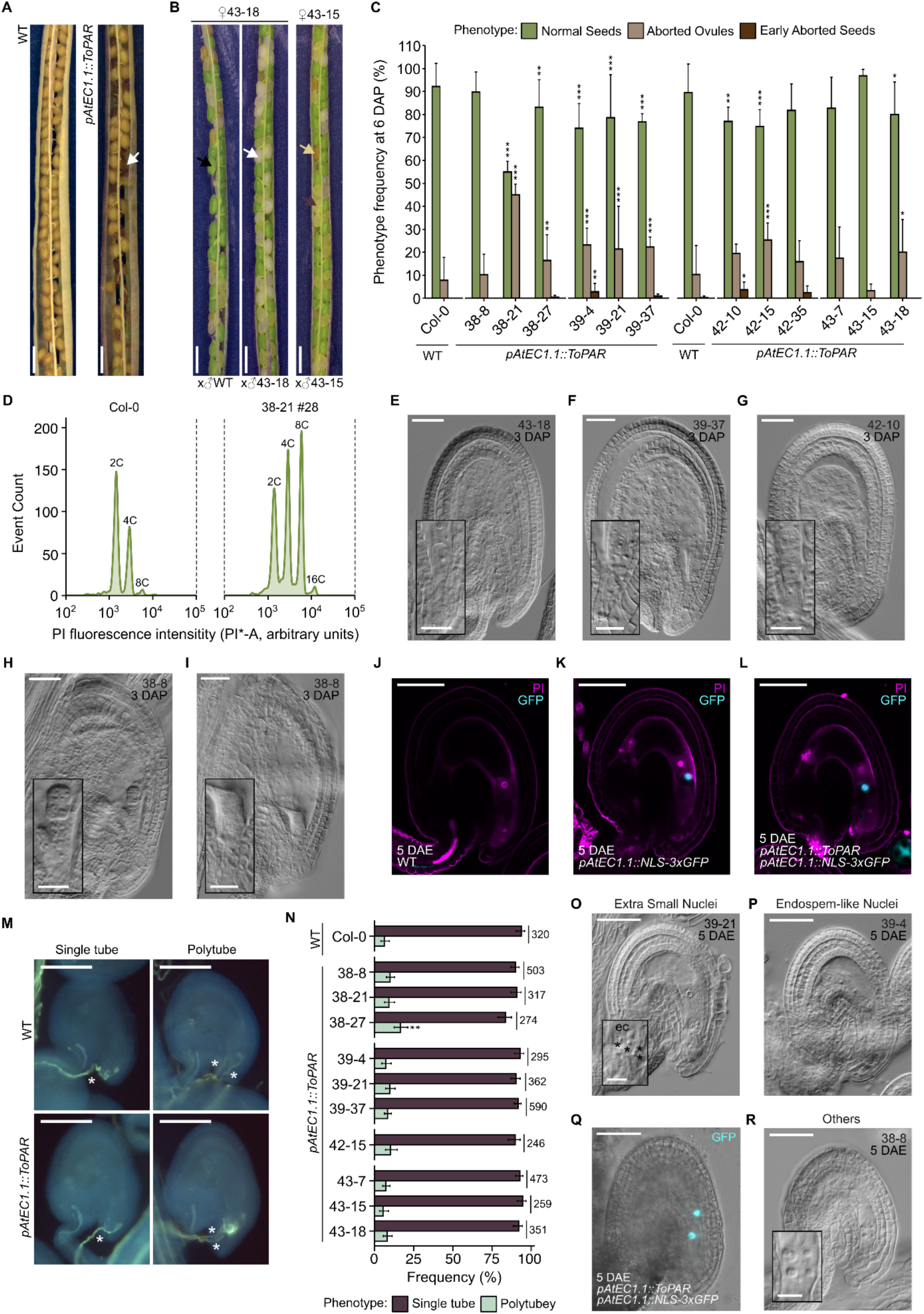
*ToPAR* expression in egg cells causes embryo sac aberrant divisions and impacts plant fertility. (**A**) Representative photo of dry WT (left) and *pAtEC1.1::ToPAR* (right) siliques. White arrow points to an aborted seed. Scale bars, 1 mm. (**B**) Siliques from lines 43-18 and 43-15 crossed with WT and/or selfed at 12 days after pollination (DAP). Black, yellow, purple and white arrows denote green, aborted, pale and white seeds, respectively. Scale bars, 1 mm. (**C**) T2 seed phenotype quantification at 6 DAP. Three siliques were quantified per genotype. Error bars represent standard deviation. Significance of differences was determined by a two-sided Fisher’s exact test. (**D**) Ploidy determination of WT and one representative *pAtEC1.1::ToPAR* T2 offspring plant showing diploid peaks. PI, propidium iodide. (**E-G**) Microscopic images of transgenic seeds with zygote (**E**), embryo with one nucleus at the embryo proper (EP) (**F**), embryo with two nuclei at the EP (**G**), abnormal embryo (**H**) and unclear phenotype (**I**) at 3 DAP. Scale bars, 50 μm. (**J-L**) Representative images of WT (**J**), *pAtEC1.1::NLS-3xGFP* (**K**) and *pAtEC1.1::ToPAR pAtEC1.1::NLS-3xGFP* (**L**) ovules at 5 DAE. Ovules were stained with PI. Scale bars, 50 μm. (**M**) Representative images of aniline blue staining of pollen tubes from crosses of WT (upper panel) and T3 *pAtEC1.1::ToPAR* (lower panel) with WT pollen. Single and multiple pollen tubes in an ovule are depicted in right and left panels, respectively. White asterisks denote pollen tubes. Scale bars, 50 μm. (**N**) Frequency of polytubey in WT and T3 *pAtEC1.1::ToPAR* ovules crossed with WT pollen at 12 hours after pollination (HAP). Three plants were analysed per line. Side values indicate the number of seeds analyzed per cross. Error bars indicate 95% Wilson confidence intervals.****p < 0.0001, ***p < 0.001, **p < 0.01 and *p < 0.05. (**O**) *pAtEC1.1::ToPAR* ovule with extra small nuclei nearby the egg cell at 5 days after emasculation (5 DAE).Scale bar, 50 μm. Asterisks denote nuclei in the egg cell apparatus. (**P**) Transgenic ovule with an endosperm-like phenotype at 5 DAE. Scale bar, 50 μm. (**Q**) *pAtEC1.1::ToPAR pAtEC1.1::NLS-3xGFP* ovule with two nuclei showing egg cell identity. Scale bar, 50 μm. (**R**) Example of a 5 DAE transgenic ovule with a phenotype classified as “Others”. Scale bars, 50 μm (12.5 μm for inset).

**Figure S3.**
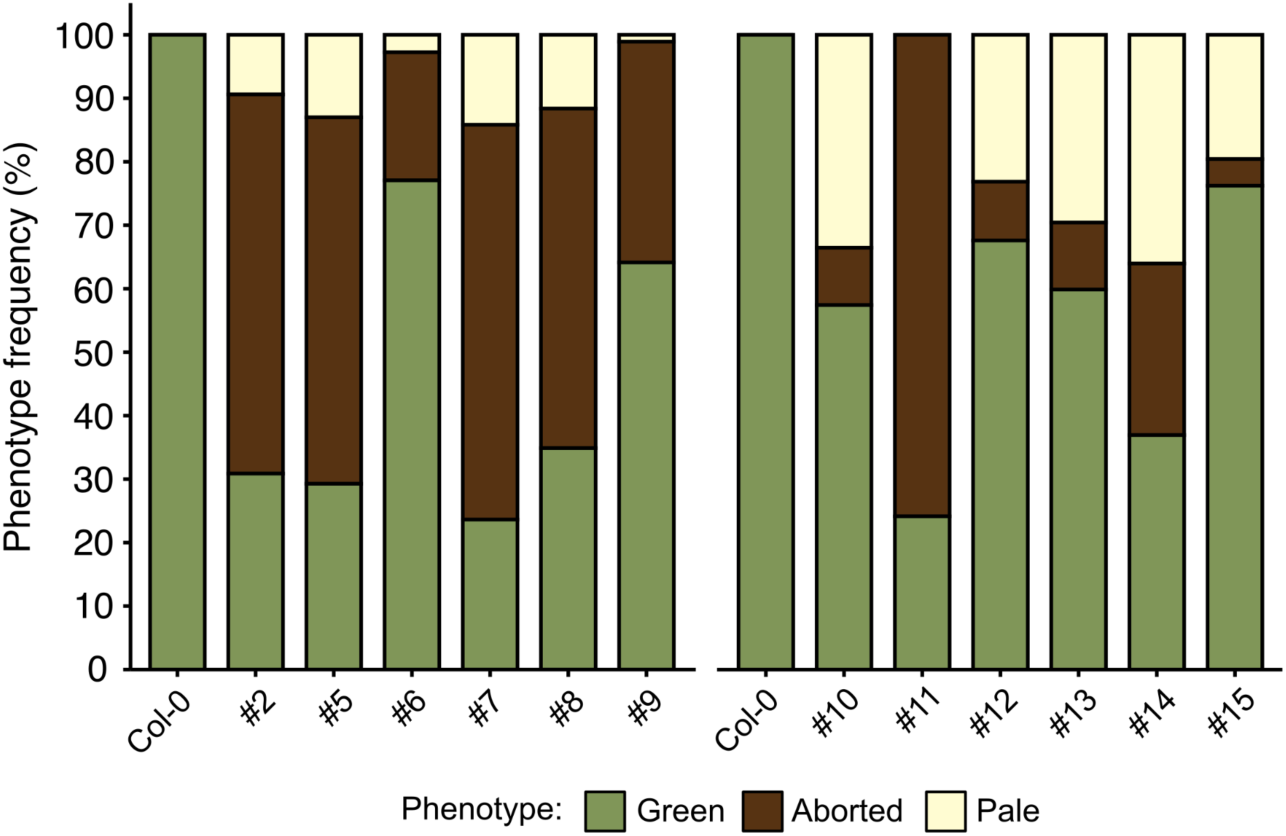
*pAtWOX8::ToPAR* T1 seed phenotype screening. Phenotype frequencies of additional T1 independent lines. Lines 1, 3 and 4 were selected for further analysis and T1 quantifications are shown in **Figure S4A**. Three siliques were quantified per independent line.

**Figure S4.**
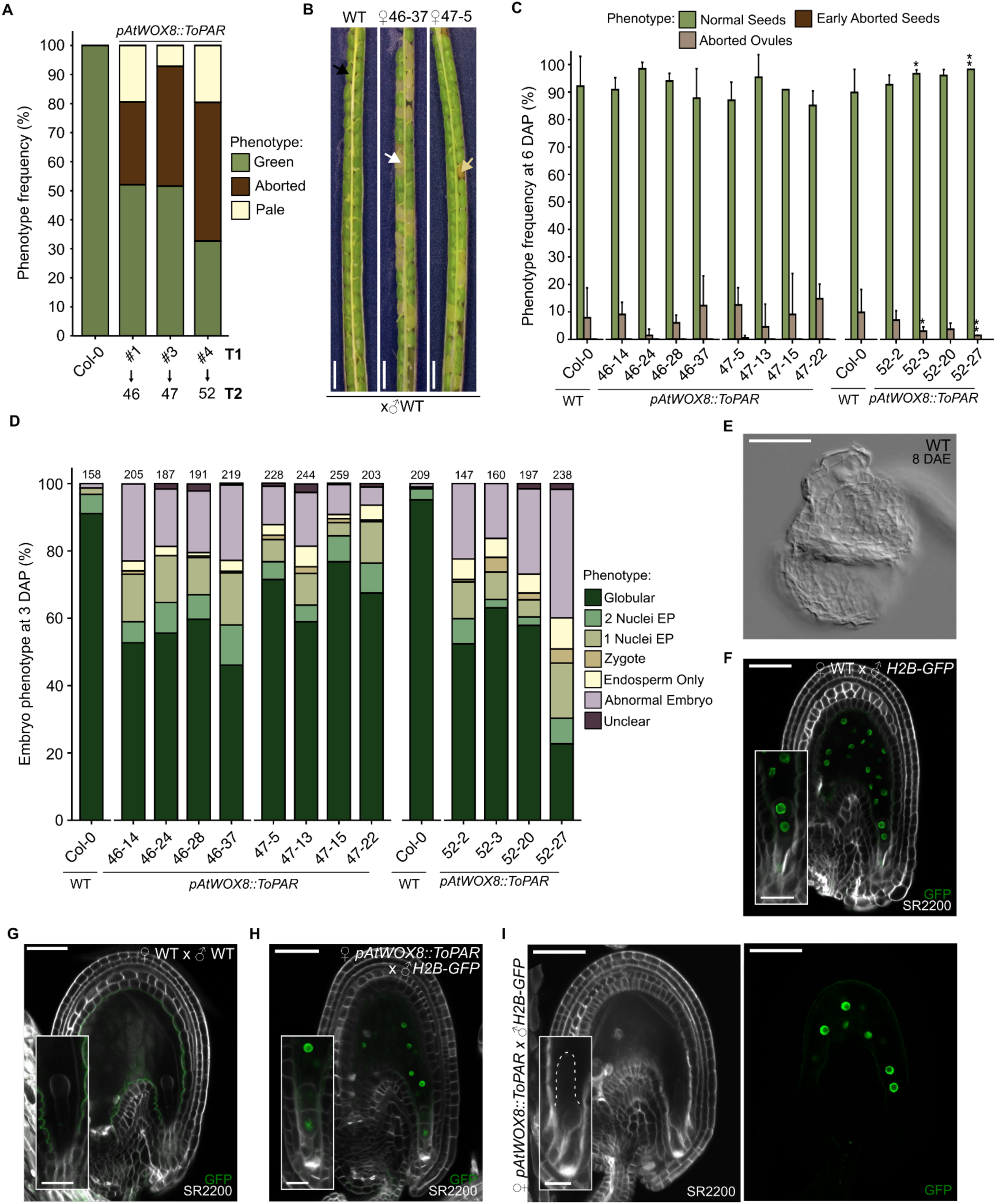
*ToPAR* expression reduces plant fertility by inducing abnormal embryogenesis. (**A**) T1 seed phenotype quantification at 12 days after pollination (DAP). (**B**) WT and *pAtWOX8::ToPAR* T2 siliques at 12 DAP. Black, white and yellow arrows denote green, pale and aborted seeds, respectively. Scale bars, 50 μm. (**C**) T2 seed phenotype quantification at 6 DAP. Three siliques were quantified per genotype. Error bars represent standard deviation. Significance of differences was determined by a two-sided Fisher’s exact test. (**D**) *pAtWOX8::ToPAR* T2 embryo phenotype frequency at 3 DAP. Top values indicate the number of seeds analyzed per genotype. (**E**) Aborted WT ovule at 8 days after emasculation (DAE). Scale bar, 50 μm. (**F-G**) Confocal images of WT ovules pollinated with *pRPS5A::H2B-GFP* (*H2B-GFP*) (**F**) and WT (**G**) at 2 DAP. Scale bars, 50 μm (25 μm for insets). (**H**) *pAtWOX8::ToPAR* seed with an embryo with a putative first symmetric zygote division. Scale bar, 50 μm (25 μm for inset). (**I**) SR2200 (left) and GFP (right) channels of a *pAtWOX8::ToPAR* ovule pollinated with *H2B-GFP* showing absent GFP signal in the micropylar region at 2 DAP. The dashed line outlines the embryo. Scale bar, 50 μm (25 μm for inset). Seeds were stained with SR2200 (in white). ****p < 0.0001, ***p < 0.001, **p < 0.01 and *p < 0.05.

**Table S1.**
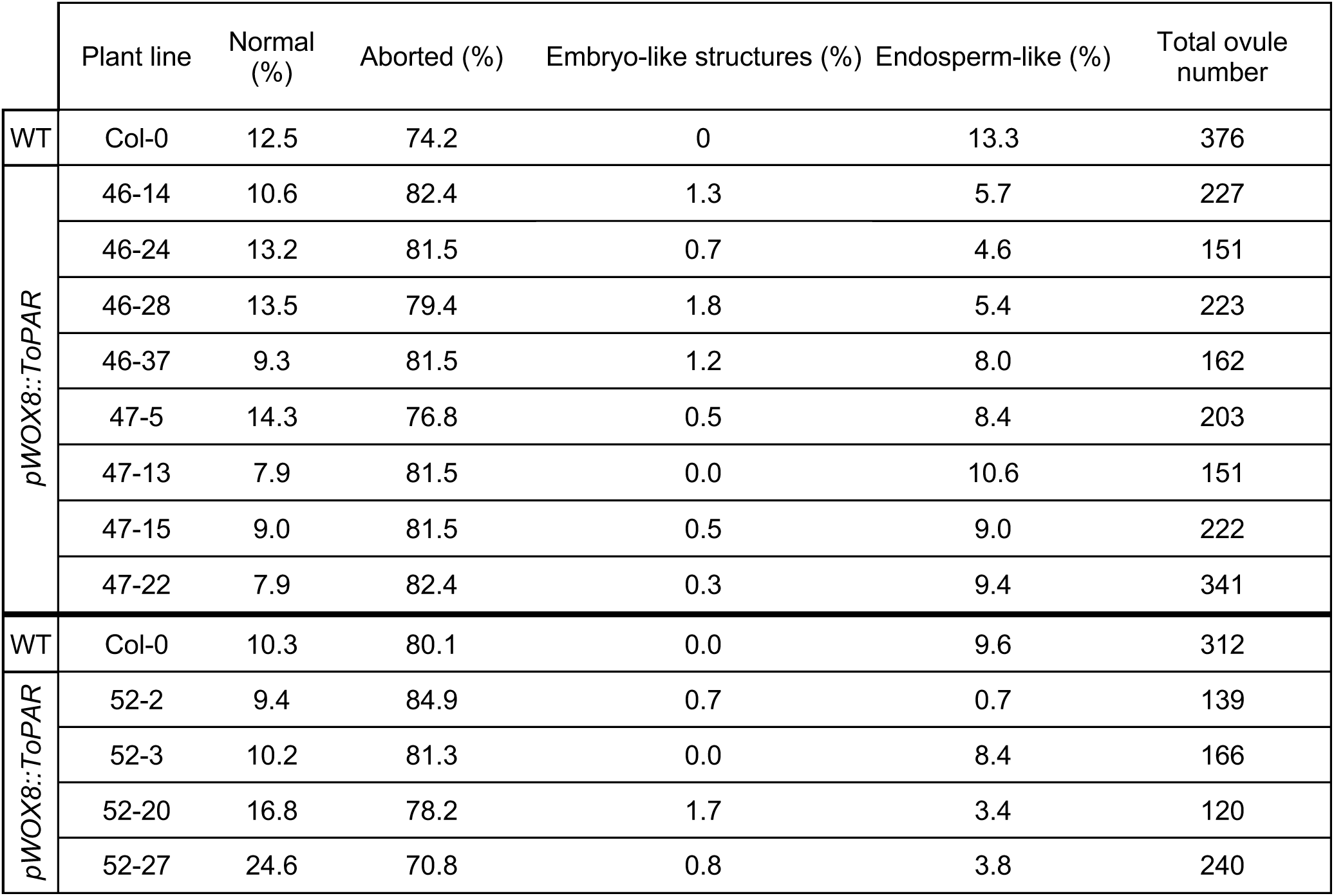
Ovule phenotype quantifications at 8 days after emasculation (DAE).

**Table S2.**
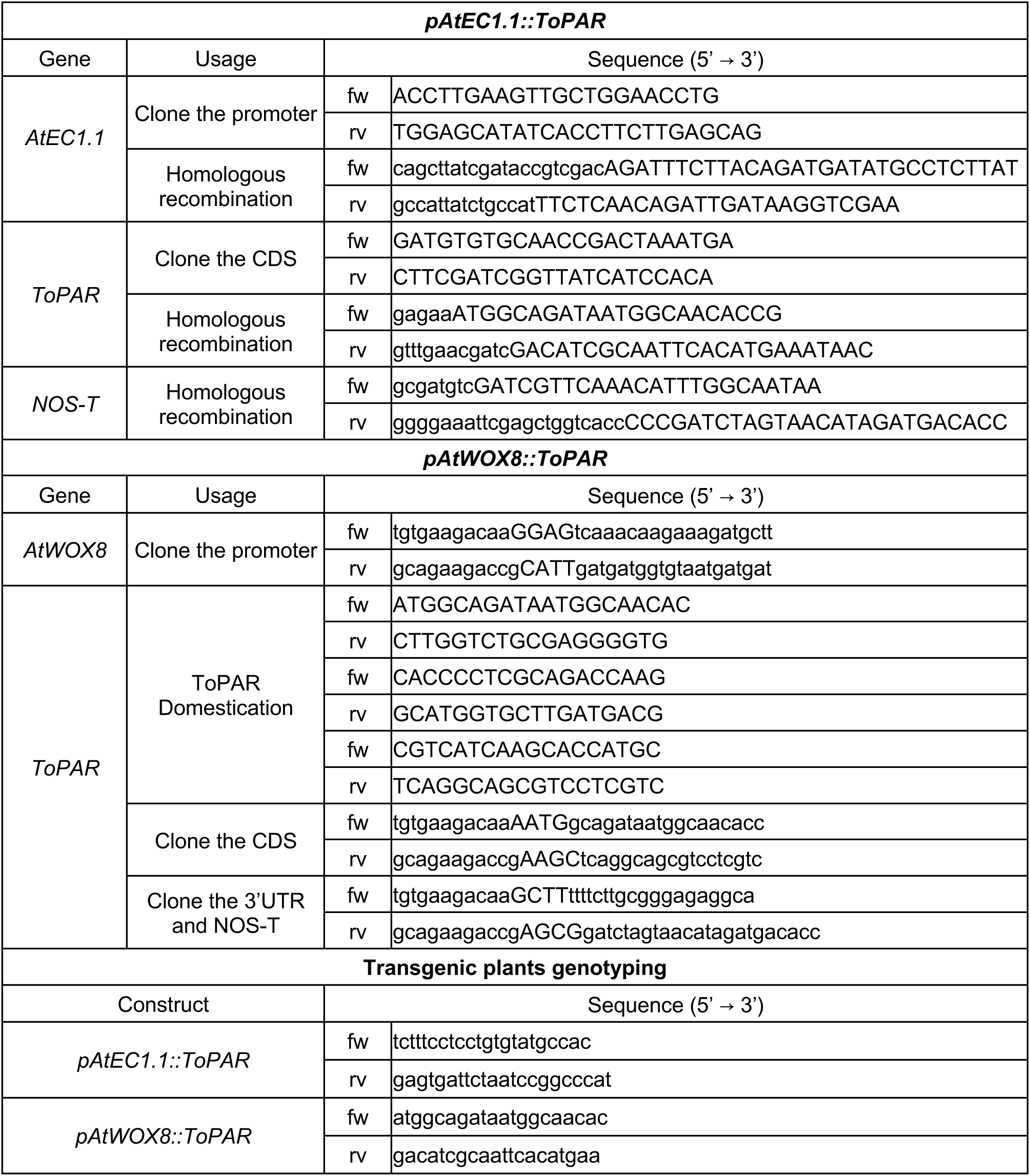
Primer sequences list.

