## Supplementary figures and tables for "The dandelion *PARTHENOGENESIS* gene dominantly modifies *Arabidopsis* fertilization and embryogenesis"

**Figure S1. *pAtEC1.1::ToPAR* T1 seed phenotype screening.**

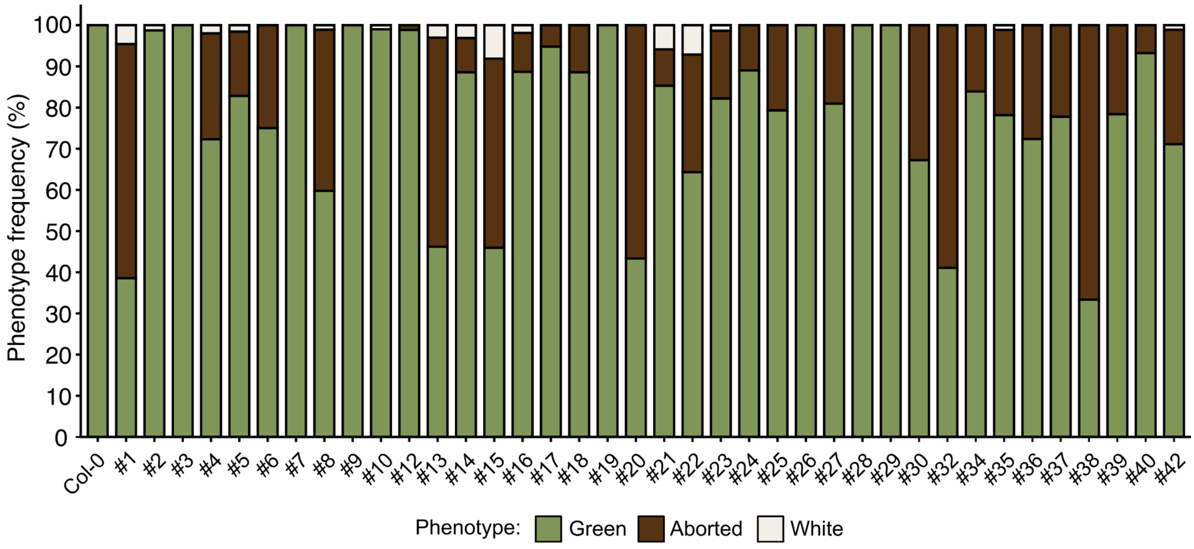


Phenotype frequencies of additional T1 independent lines. Two-three siliques were quantified per independent line.

**Figure S2. *ToPAR* expression in egg cells causes embryo sac aberrant divisions and impacts plant fertility.**

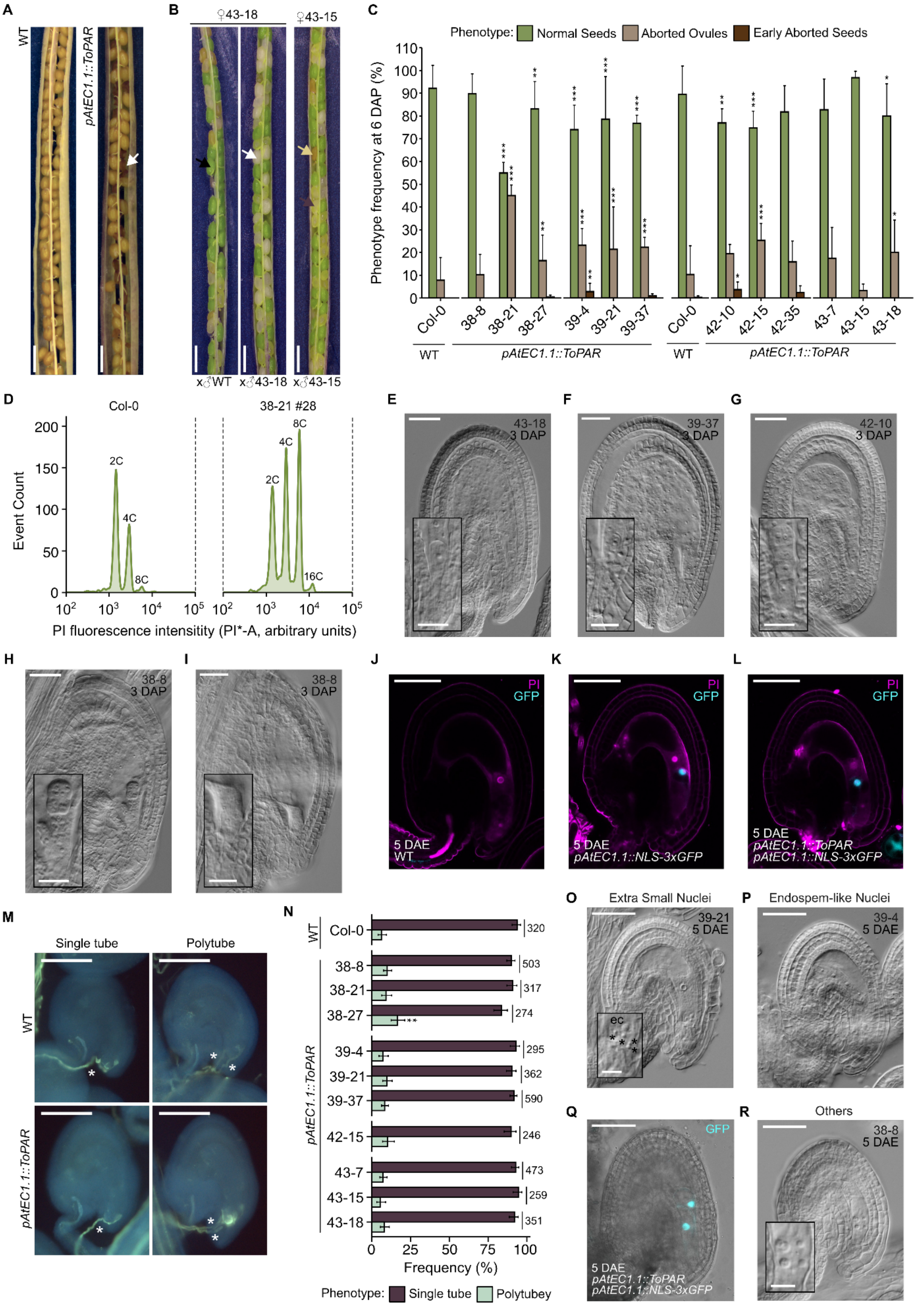

**Figure S2.** ***ToPAR*** **expression in egg cells causes embryo sac aberrant divisions and impacts plant fertility.**

(**A**) Representative photo of dry WT (left) and *pAtEC1.1::ToPAR* (right) siliques. White arrow points to an aborted seed. Scale bars, 1 mm. (**B**) Siliques from lines 43-18 and 43-15 crossed with WT and/or selfed at 12 days after pollination (DAP). Black, yellow, purple and white arrows denote green, aborted, pale and white seeds, respectively. Scale bars, 1 mm. (**C**) T2 seed phenotype quantification at 6 DAP. Three siliques were quantified per genotype. Error bars represent standard deviation. Significance of differences was determined by a two-sided Fisher's exact test. (**D**) Ploidy determination of WT and one representative *pAtEC1.1::ToPAR* T2 offspring plant showing diploid peaks. PI, propidium iodide. (**E-G**) Microscopic images of transgenic seeds with zygote (**E**), embryo with one nucleus at the embryo proper (EP) (**F**), embryo with two nuclei at the EP (**G**), abnormal embryo (**H**) and unclear phenotype (**I**) at 3 DAP. Scale bars, 50 µm. (**J-L**) Representative images of WT (**J**), *pAtEC1.1::NLS-3xGFP* (**K**) and *pAtEC1.1::ToPAR pAtEC1.1::NLS-3xGFP* (**L**) ovules at 5 DAE. Ovules were stained with PI. Scale bars, 50 µm. (**M**) Representative images of aniline blue staining of pollen tubes from crosses of WT (upper panel) and T3 *pAtEC1.1::ToPAR* (lower panel) with WT pollen. Single and multiple pollen tubes in an ovule are depicted in right and left panels, respectively. White asterisks denote pollen tubes. Scale bars, 50 µm. (**N**) Frequency of polytubey in WT and T3 *pAtEC1.1::ToPAR* ovules crossed with WT pollen at 12 hours after pollination (HAP). Three plants were analysed per line. Side values indicate the number of seeds analyzed per cross. Error bars indicate 95% Wilson confidence intervals.****$p < 0.0001$, ***$p < 0.001$, **$p < 0.01$ and *$p < 0.05$. (**O**) *pAtEC1.1::ToPAR* ovule with extra small nuclei nearby the egg cell at 5 days after emasculation (5 DAE).Scale bar, 50 µm. Asterisks denote nuclei in the egg cell apparatus. (**P**) Transgenic ovule with an endosperm-like phenotype at 5 DAE. Scale bar, 50 µm. (**Q**) *pAtEC1.1::ToPAR pAtEC1.1::NLS-3xGFP* ovule with two nuclei showing egg cell identity. Scale bar, 50 µm. (**R**) Example of a 5 DAE transgenic ovule with a phenotype classified as "Others". Scale bars, 50 µm (12.5 µm for inset).

**Figure S3. *pAtWOX8::ToPAR* T1 seed phenotype screening.**

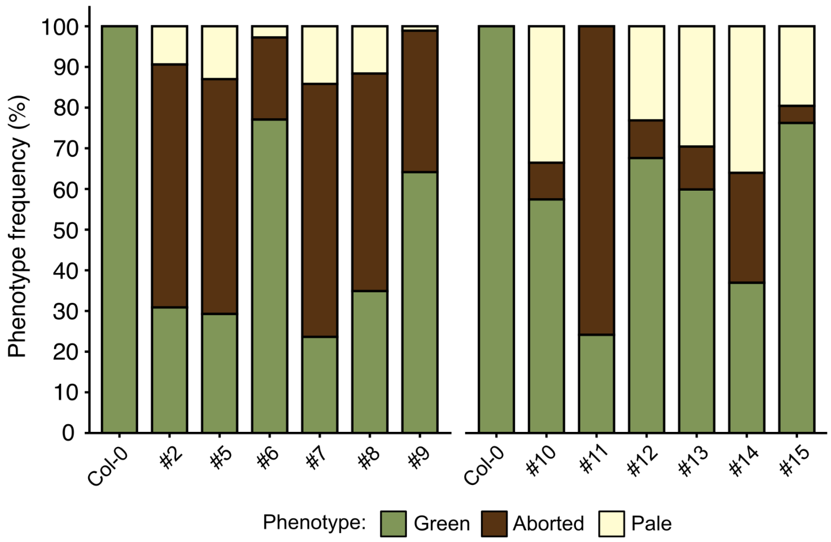


Phenotype frequencies of additional T1 independent lines. Lines 1, 3 and 4 were selected for further analysis and T1 quantifications are shown in **Figure S4A**. Three siliques were quantified per independent line.

**Figure S4. *ToPAR* expression reduces plant fertility by inducing abnormal embryogenesis.**

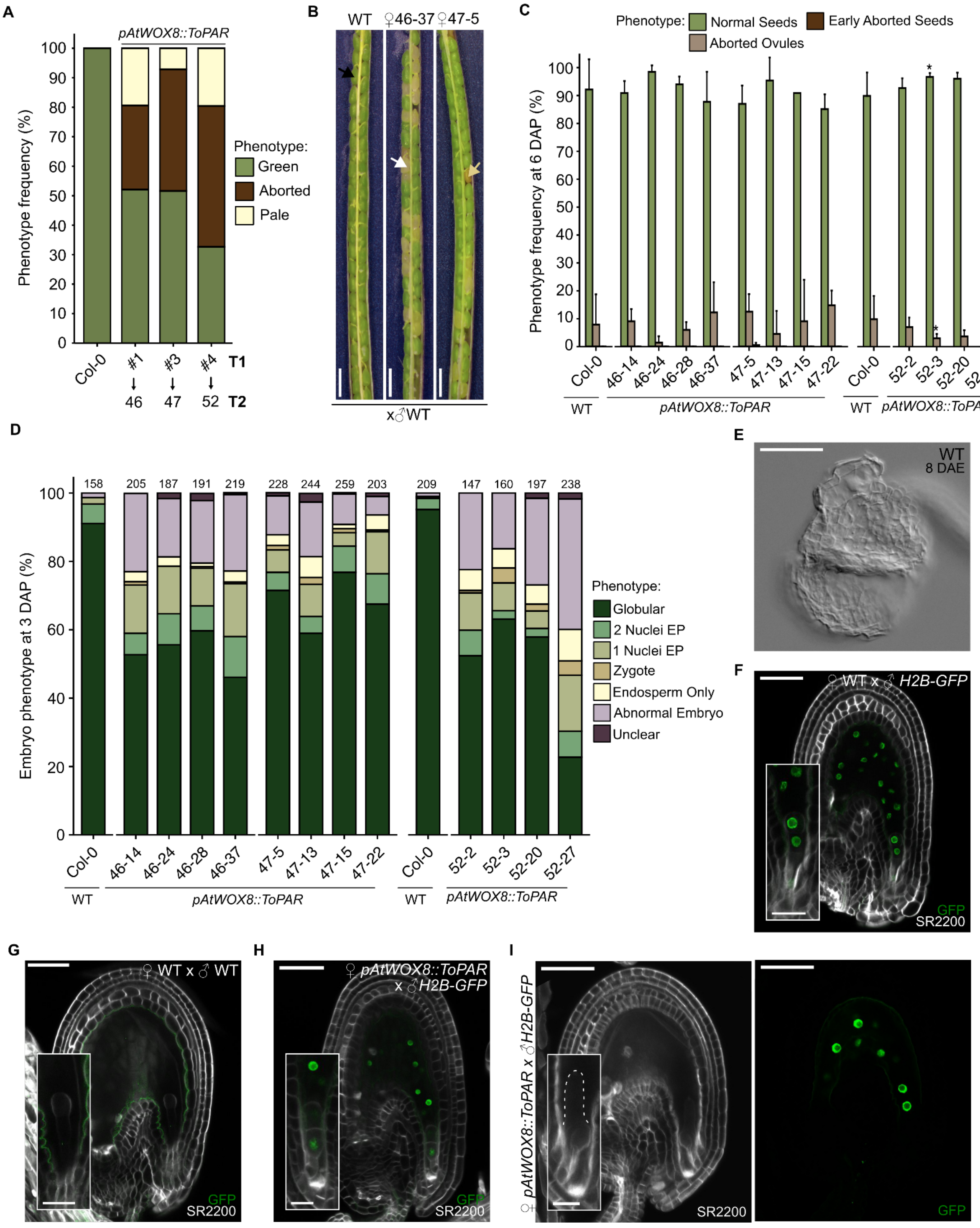

**Figure S4. *ToPAR* expression reduces plant fertility by inducing abnormal embryogenesis.**

(**A**) T1 seed phenotype quantification at 12 days after pollination (DAP). (**B**) WT and *pAtWOX8::ToPAR* T2 siliques at 12 DAP. Black, white and yellow arrows denote green, pale and aborted seeds, respectively. Scale bars, 50 µm. (**C**) T2 seed phenotype quantification at 6 DAP. Three siliques were quantified per genotype. Error bars represent standard deviation. Significance of differences was determined by a two-sided Fisher's exact test. (**D**) *pAtWOX8::ToPAR* T2 embryo phenotype frequency at 3 DAP. Top values indicate the number of seeds analyzed per genotype. (**E**) Aborted WT ovule at 8 days after emasculation (DAE). Scale bar, 50 µm. (**F-G**) Confocal images of WT ovules pollinated with *pRPS5A::H2B-GFP* (*H2B-GFP*) (**F**) and WT (**G**) at 2 DAP. Scale bars, 50 µm (25 µm for insets). (**H**) *pAtWOX8::ToPAR* seed with an embryo with a putative first symmetric zygote division. Scale bar, 50 µm (25 µm for inset). (**I**) SR2200 (left) and GFP (right) channels of a *pAtWOX8::ToPAR* ovule pollinated with *H2B-GFP* showing absent GFP signal in the micropylar region at 2 DAP. The dashed line outlines the embryo. Scale bar, 50 µm (25 µm for inset). Seeds were stained with SR2200 (in white). ****$p < 0.0001$, ***$p < 0.001$, **$p < 0.01$ and *$p < 0.05$.

**Table S1. Ovule phenotype quantifications at 8 days after emasculation (DAE).**

| | Plant line | Normal (%) | Aborted (%) | Embryo-like structures (%) | Endosperm-like (%) | Total ovule number |
|---|---|---|---|---|---|---|
| WT | Col-0 | 12.5 | 74.2 | 0 | 13.3 | 376 |
| *pWOX8::ToPAR* | 46-14 | 10.6 | 82.4 | 1.3 | 5.7 | 227 |
| | 46-24 | 13.2 | 81.5 | 0.7 | 4.6 | 151 |
| | 46-28 | 13.5 | 79.4 | 1.8 | 5.4 | 223 |
| | 46-37 | 9.3 | 81.5 | 1.2 | 8.0 | 162 |
| | 47-5 | 14.3 | 76.8 | 0.5 | 8.4 | 203 |
| | 47-13 | 7.9 | 81.5 | 0.0 | 10.6 | 151 |
| | 47-15 | 9.0 | 81.5 | 0.5 | 9.0 | 222 |
| | 47-22 | 7.9 | 82.4 | 0.3 | 9.4 | 341 |
| WT | Col-0 | 10.3 | 80.1 | 0.0 | 9.6 | 312 |
| *pWOX8::ToPAR* | 52-2 | 9.4 | 84.9 | 0.7 | 0.7 | 139 |
| | 52-3 | 10.2 | 81.3 | 0.0 | 8.4 | 166 |
| | 52-20 | 16.8 | 78.2 | 1.7 | 3.4 | 120 |
| | 52-27 | 24.6 | 70.8 | 0.8 | 3.8 | 240 |

**Table S2. Primer sequences list.**

| ***pAtEC1.1::ToPAR*** | | | |
|---|---|---|---|
| Gene | Usage | | Sequence (5' → 3') |
| *AtEC1.1* | Clone the promoter | fw | ACCTTGAAGTTGCTGGAACCTG |
| | | rv | TGGAGCATATCACCTTCTTGAGCAG |
| | Homologous recombination | fw | cagcttatcgataccgtcgacAGATTTCTTACAGATGATATGCCTCTTAT |
| | | rv | gccattatctgccatTTCTCAACAGATTGATAAGGTCGAA |
| *ToPAR* | Clone the CDS | fw | GATGTGTGCAACCGACTAAATGA |
| | | rv | CTTCGATCGGTTATCATCCACA |
| | Homologous recombination | fw | gagaaATGGCAGATAATGGCAACACCG |
| | | rv | gtttgaacgatcGACATCGCAATTCACATGAAATAAC |
| *NOS-T* | Homologous recombination | fw | gcgatgtcGATCGTTCAAACATTTGGCAATAA |
| | | rv | ggggaaattcgagctggtcaccCCCGATCTAGTAACATAGATGACACC |
| ***pAtWOX8::ToPAR*** | | | |
| Gene | Usage | | Sequence (5' → 3') |
| *AtWOX8* | Clone the promoter | fw | tgtgaagacaaGGAGtcaaacaagaaagatgctt |
| | | rv | gcagaagaccgCATTgatgatggtgtaatgatgat |
| *ToPAR* | ToPAR Domestication | fw | ATGGCAGATAATGGCAACAC |
| | | rv | CTTGGTCTGCGAGGGGTG |
| | | fw | CACCCCTCGCAGACCAAG |
| | | rv | GCATGGTGCTTGATGACG |
| | | fw | CGTCATCAAGCACCATGC |
| | | rv | TCAGGCAGCGTCCTCGTC |
| | Clone the CDS | fw | tgtgaagacaaAATGgcagataatggcaacacc |
| | | rv | gcagaagaccgAAGCtcaggcagcgtcctcgtc |
| | Clone the 3'UTR and NOS-T | fw | tgtgaagacaaGCTTttttcttgcgggagaggca |
| | | rv | gcagaagaccgAGCGgatctagtaacatagatgacacc |
| **Transgenic plants genotyping** | | | |
| Construct | | | Sequence (5' → 3') |
| *pAtEC1.1::ToPAR* | | fw | tctttcctcctgtgtatgccac |
| | | rv | gagtgattctaatccggcccat |
| *pAtWOX8::ToPAR* | | fw | atggcagataatggcaacac |
| | | rv | gacatcgcaattcacatgaa |
